# Design of Worst-Case-Optimal Spaced Seeds

**DOI:** 10.1101/2023.11.20.567826

**Authors:** Sven Rahmann, Jens Zentgraf

## Abstract

Read mapping (and alignment) is a fundamental problem in biological sequence analysis. For speed and computational efficiency, many popular read mappers tolerate only a few differences between the read and the corresponding part of the reference genome, which leads to reference bias: Reads with too many difference are not guaranteed to be mapped correctly or at all, because to even consider a genomic position, a sufficiently long *exact* match (*seed*) must exist.

While pangenomes and their graph-based representations provide one way to avoid reference bias by enlarging the reference, we explore an orthogonal approach and consider stronger substitution-tolerant primitives, namely *spaced seeds* or gapped *k*-mers. Given two integers *k ≤ w*, one considers *k* selected positions, described by a *mask*, from each length-*w* window in a sequence. In the existing literature, masks with certain *probabilistic* guarantees have been designed for small values of *k*.

Here, for the first time, we take a combinatorial approach from a *worst-case* perspective. For any mask, using integer linear programs, we find least favorable distributions of sequence changes in two different senses: (1) minimizing the number of unchanged windows; (2) minimizing the number of positions covered by unchanged windows. Then, among all masks of a given shape (*k, w*), we find the set of best masks that maximize these minima. As a result, we obtain highly robust masks, even for large numbers of changes. Their advantages are illustrated in two ways: First, we provide a new challenge dataset of simulated DNA reads, on which current methods like bwa-mem2, minimap2, or strobealign struggle to find seeds, and therefore cannot produce alignments against the human t2t reference genome, whereas we are able to find the correct location from a few unique spaced seeds. Second, we use real DNA data from the highly diverse human HLA region, which we are able to map correctly based on a few exactly matching spaced seeds of well-chosen masks, without evaluating alignments.

## 1 Introduction

Read mapping (and alignment) is a fundamental problem in biological sequence analysis required for many tasks in genomics and transcriptomics, like genome-wide variant calling, transcript expression quantification, or species identification and quantification in metagenomics, to name just a few. Today, there exist efficient methods (in terms of both memory and running time) to map millions of reads against a reference genome, such as bwa-mem2 [20], bowtie2 [10], minimap2 [11, 12], strobealign [19] and others.

Read mappers must balance efficiency against error-tolerance: Finding *exact* matches between reads and reference is extremely efficient with the existing index data structures, but allowing an increasing number of differences (typically measured in terms of edit distance) between a read and the reference requires time that increases exponentially with the number of tolerated differences, because one has to evaluate many branching paths in an index data structure [13]. Thus, for efficiency, the default settings of popular read mappers allow only a few erros, which leads to *reference bias*: Reads that are too different from their best match in the reference genome may not be mapped at all, and the corresponding genomic variants may be missed during downstream variant calling steps [14].

One answer to the problem of reference bias has been the construction of pangenomes that represent not only a single linear sequence, but a large collection of possible genome sequences [4] of a species that can be compactly represented by a branching graph, such that known sequence variants are included in the pangenome. Consequently, read mapping algorithms and their underlying index data structures, have been generalized to enable mapping linear reads against graph reference genomes [6, 18, 15]. Because many of the known variants are included in the reference, a small error threshold is now sufficient where a possibly much larger one was required before.

An orthogoal approach is to design index data structures with built-in error tolerance. This appears to be much easier for Hamming distance (substitutions only) than for general edit distance (substitutions, insertions and deletions), because of the variability of the length of the match when indels are allowed. In the past, *gapped k-mers*, also known as *spaced seeds*, have been proposed and used as substitution-tolerant primitives: One only considers *k* specifically selected positions, described by a *mask*, out of a larger window of *w≥ k* positions. There is a rich body of literature on finding masks for given (*k, w*) perform well or even optimally in a specific sense [1, 2, 3, 5, 8, 9, 16, 23].

Most of this exising work has been of a probabilistic nature, i.e., about computing or optimizing the probability that there is at least one matching seed (“hit”) under different assumptions about sequence similarity. The optimal mask(s) unsurprisingly depend on the model parameters (evolutionary time, expected number of changed basepairs), but at least for simple Bernoulli models, where a change can appear at each sequence position independently with some small probability *p*, the problem has been comprehensively solved: The probability of at least one hit can computed as parameterized polynomial in *p*, from which one can identify the small set of masks that are optimal for some value of *p*, or integrated over a certain *p*-interval. In essence, one uses dynamic programming to count (or accumulate probabilities of) binary sequences that do not contain the mask as a substring; these calculations can be carried out symbolically.

We refer to the comprehensive paper of Laurent Noé [16] for details. These methods are computationally expensive; so most experiments were limited to small values of *k*, i.e. *k* = 11, *w ≤* 22, and short sequences of length *n≤* 64.

Perhaps surprisingly, the mask choice problem (or spaced seed design problem) has never been approached completely from a worst-case perspective. In this work, we close this open gap by evaluating masks based on their worst-case performance (for a given sequence length and a fixed number of changes) by considering the following two combinatorial optimization problems.

**MinHits** For a given sequence length *n* (a typical short read, or part of a longer read) and a given number *c* of allowed sequence changes (substitutions only), and a given spaced seed pattern (also called mask), find the worst-case distribution of the *c* changes that minimizes the number of seed *hits* inside the sequence (formal definitions below).

**MinCov** Under the same assumptions, find the distribution of the *c* changes that minimizes the number of sequence positions covered by the hits. (This is an alternative notion of “worst-case”.)

Solving the above two problems for a given mask yields its worst-case change distributions. This means that the resulting number of hits (resp. covered positions) are guaranteed for this mask for *every* distribution of *c* changes across the sequence. Of course, if the number of changes *c* is chosen too large, the objective value for both problems is zero, so there is a limited range of interest for parameter *c*. The most interesting problem, however, is to find a “best” mask where these minima are maximal (in a specified class of masks).

**MaxMinHits** Given a sequence length *n*, a number of changes *c*, and a set ℳ of masks, find the masks that maximize the MinHits objective among all masks in ℳ.

**MaxMinCov** Under the same assumptions, find the masks in that maximize the MinCov objective in ℳ.

We shall now define the necessary preliminaries and provide an example; then we give formal definitions of the above problems. We solve them for practically relevant seed parameters (*k, w*), sequence lengths *n* and numbers of changes *c*. As a result, we obtain highly substitution-tolerant seeds with guarantees that do not only hold with high probability but always. We illustrate the impact of using such seeds on both a simulated challenge dataset and real MiSeq data from the diverse human HLA region.

## 2 Methods

We start by defining the necessary terms to work with spaced seeds, or gapped *k*-mers; in particular we define *masks* of a given *shape* (*k, w*). Then, we introduce the optimization problems of computing the worst-case set of change positions, and the problem of finding the best mask among all masks of shape (*k, w*). These problems are formulated as integer linear programs (ILPs). In addition, we introduce the notion of (strongly) unique gapped *k*-mers in a reference genome.

### 2.1 Definitions

#### Basics

We write [*n*] for the set of integers *{*0, …, *n* − 1*}*. Indexing of strings starts at 0, so *s* = (*s*_0_, …, *s*_*n*−1_) = (*s*_*i*_)_*i*∈[*n*]_, where the length of *s* is |*s*| = *n*.

We consistently use a single (reference) sequence *s* in this manuscript; however, multiple sequences (e.g., chromosomes) are covered by concatenating them with separator characters $, ignoring all (gapped) *k*-mers containing a separator, and translating between single sequence positions and chromosome-position-pairs accordingly.

If *s* and *t* are strings of equal length |*s*| = |*t*| = *n*, then their *Hamming distance d*_H_(*s, t*) is the number of indices *i* ∈ [*n*] where *s*_*i*_*≠*= *t*_*i*_.

#### Masks of shape (*k, w*)

Given two integers *w≥ k ≥* 2, a (*k, w*)-*mask* is a string *μ* of length *w* over the alphabet {`#, _`} that contains exactly *k* times the character `#` and *w*− *k* times the character `_`. The positions marked `#` are called *significant*, and the positions marked _ are called *insignificant* or *gaps*. We call *k* the *weight* of the mask and *w* its *width* or *window length*. The pair (*k, w*) is called the *shape* of the mask. A mask *μ* may also be represented as the tuple *κ* of significant positions: *κ* = *{j* : 0 *≤ j < w* and *μ*_*j*_ = `#`*}*.

We require that *μ*_0_ = *μ*_*w*−1_ = `#`, because otherwise the mask could be shortened by removing the insignificant characters at each end. With this constraint, there are 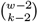 different masks of shape (*k, w*).

A mask *μ* is *symmetric* if *μ* = *rev*(*μ*) := *μ* _*n*−1_ … *μ*_1_*μ*_0_. We consider only symmetric masks in this work because then we can work with canonical gapped *k*-mers, as the mask is the same on both DNA strands. We also restrict ourselves to odd *k* and *w* to avoid any issues with *k*-mers that are equal to their reverse complement. For symmetric masks with odd *k* and *w*, we must have *μ* = `#` for the middle position *m* = (*w* − 1)*/*2, and there are ^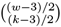^ such masks. They can be enumerated by enumerating all possible left halves and mirroring them to obtain the right halves.

We write ℳ (*k, w*) for the set of all mask tuples *κ* (with significant ends) of shape (*k, w*) and *S*(*k, w*) for the subset of all symmetric ones. For example, *κ* = (0, 2, 5, 8, 10) ∈ 𝒮 (5, 11), corresponding to *μ* = `#_# # #_#`. For *k* = 25 and *w* = 37, we find | ℳ (*k, w*)| = 834 451 800 and | 𝒮 (*k, w*)| = 12 376. It becomes clear that symmetry is a strong constaint.

#### *κ*-mers of a sequence

Given a mask *μ* of shape (*k, w*) with its corresponding offset tuple *κ* and a string *s* of length *n* over an alphabet (e.g., the DNA or protein alphabet), we obtain *n* − *w* + 1 *gapped k-mers* from *s*: The *i*-th gapped *k*-mer *g*_*i*_, for *i* ∈ [*n* − *w*], is obtained by concatenating the significant positions when the mask is applied at positions *i*, …, *i* + *w* −1 of *s*, i.e., *g*_*i*_ = (*s*_*i*+*j*_)_*j*∈*κ*_. We also say that each such string *g*_*i*_ is a *μ-mer* or *κ-mer* of *s*, depending on whether we are referring to the mask’s string representation *μ* or tuple representation *κ*. In this sense, a (contiguous) *k*-mer of *s* is a *κ* = (0, 1, …, *k* − 1)-mer of *s*.

#### Effects of sequence changes on *κ*-mers

When we change a single character in a sequence *s*, then up to *k* of the *κ*-mers may be changed (less if the changed position is near either end of the sequence). We say that the *κ*-mer starting at position *p*∈ [*n − w* + 1] (*κ*-mer number *p* for short) is *changed* or *destroyed* if any of the positions *p* + *j* : *j* ∈ *κ* is changed. Otherwise, the *κ*-mer is *unchanged* or *unaffected* ; we also say that it *hits* the sequence.

We say that sequence position *i* ∈ [*n*] is *covered* (by an unaffected *κ*-mer) if there exists an unaffected *κ*-mer starting at some position *p ≤ i* such that *i* = *p* + *j* for some *j* ∈ *κ*. In other words, if *κ*-mer number *p* is unaffected, then all positions *p* + *j* for *j* ∈ *κ* are covered.

### 2.3 Optimization Problems

We introduce the different optimization problems that we consider in this work.

For a given mask *κ* of shape (*k, w*) and given sequence length *n*, it asks how many changes *C*^∗^ = *C*^∗^(*κ, n*) can be made at most in order to guarantee at least one unchanged *κ*-mer (for all possible placements of *c* changes among *n* positions). Equivalently, we may find the smallest *C*^*†*^ such that *C*^*†*^ changes, placed conspiratively, change all *κ*-mers; then *C*^∗^ = *C*^*†*^− 1. Masks of the same shape (*k, w*) may have different *C*^∗^-values (Figure 1), but even masks with the same (highest) *C*^∗^-value can be further distinguished.

**Figure 1:**
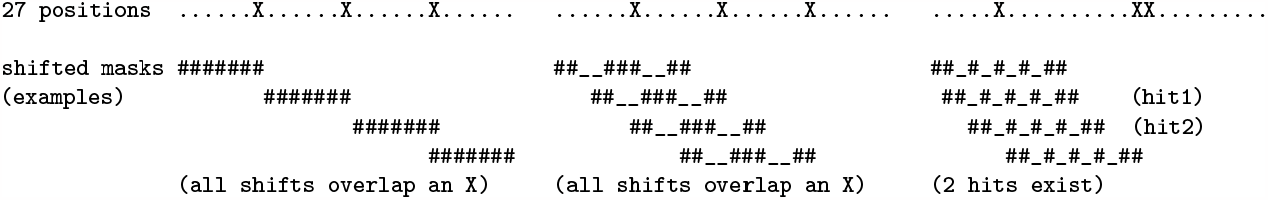
Examples for weight *k* = 7, *c* = 3 changes and sequence length *n* = 27. For contiguous 7-mers (*k* = *w* = 7, mask `#######`, left), we can place 3 changes at (0-based) positions *X* = {6, 13, 20} to change all 7-mers; so *C*^*†*^ = 3 and *C*^∗^ = 2. For mask `##--###--##` (middle) with (*k, w*) = (7, 11), we have the same result with the same placement of changes. (middle)However, using the (7, 11)-mask `##_#_#_#_##` (right), we tolerate *C*^∗^ = 3 changes and always guarantee 2 hits (unchanged gapped *k*-mers) and at least 10 covered positions. The rightmost example shows one worst-case positioning of the 3 changes to minimize hits; it is not trivial see this by eye-balling, but is a result of the **MinHits** optimization problem in Sec. 2.2.

This leads to another optimization problem for a given mask *κ*, sequence length *n*, and number of allowed changes *c* (where *c ≤ C*^∗^(*κ, n*)): Find a worst-case positioning of the changes among the *n* sequence positions. There are two variants “MinNum” and “MinCov” of the problem: Place the changes to minimize

1. **(MinHits)** the number of (starting poisitions of) unaffected *κ*-mers (“hits”);
2. **(MinCov)** the number of covered positions.

Given a shape (*k, w*), i.e., weight *k* and window length *w*, we seek a (symmetric) mask *κ* of the given shape that *maximizes*, among all such (symmetric) masks, the minimized worst-case number of hits **(MaxMinHits)** or covered positions **(MaxMinCov)**. These are *maximin* problems, and one approach to solve them is to solve MinHits and MinCov exhaustively for all masks of the given shape, and the pick the best one.

### 2.3 Integer Linear Program Formulations

We formulate the optimization problems MinHits and MinCov as integer linear programs (ILPs). The used integer variables are binary, i.e., take only values in {0, 1} . We introduce the complete set of parameters and variables here, even though some of the problems use only a subset. The given (constant) parameters are:

- *n*: length of the sequence (typically *n* = 100, corresponding to the length of a short read);
- *c*: number of allowed changes;
- *k*: weight of the mask, number of significant positions, typically *k ≥* 21 on a mammalian genome;
- *w*: window length, *k ≤ w ≤ n*
- *κ*: *k*-tuple of offsets (0, …, *w* − 1) with *k* − 2 free entries, satisfying symmetry *i* ∈ *κ* ⇐⇒ *w* − 1 − *i* ∈ *κ*;

We use the following *binary* variables:

- *x* = (*x*_*i*_)_*i*∈[*n*]_, indicators of changed positions: *x*_*i*_ = 1 if and only if sequence position *i* is changed;
- *y* = (*y*_*p*_)_*p*∈[*n*−*w*+1]_, indicators of (starting positions of) unchanged *κ*-mers: *y*_*p*_ = 1 if and only if the *κ*-mer starting at position *p* is unchanged (i.e., none of its significant positions is changed),
- *z* = (*z*_*i*_)_*i*∈[*n*]_, indicators of covered positions: *z*_*i*_ = 1 if and only if sequence position *i* is covered by an unchanged *κ*-mer.

As seen here, we use *i* ∈ [*n*] for indexing sequence positions, *p*∈ [*n* −*w* + 1] for indexing starting positions of *κ*-mers, and *j* for an element of *κ*.

#### TolChg: Tolerated Number of Changes for a Mask

We minimize the required number of changes in order to change *all κ*-mers, i.e., at least one significant position of each *κ*-mer coincides with a changed position. The number of tolerated changes *C*^∗^(*κ, n*) is one less than the optimal objective value. We only need the binary *x*-variables (see above).

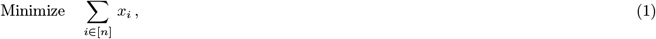

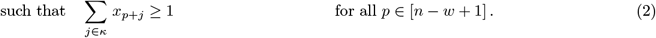

#### MinHits

Assuming that a number of changes *c ≤ C*^∗^(*κ, n*) is specified, we compute the worst-case distribution of the *c* changes that mimimizes the number of unchanged *κ*-mers. We need the *x*- and *y*-variables.

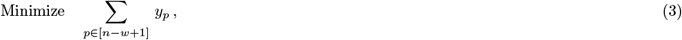

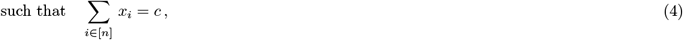

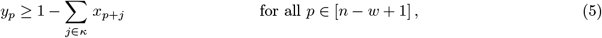

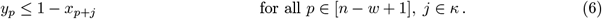

Here, Eq. (4) ensures that we use exactly *c* changes, the group of inequalities (5) ensures that *y*_*p*_ = 1 if all positions of the *κ*-mer starting at *p* are unchanged, and (6) conversely ensures that *y*_*p*_ = 0 if any position of that *κ*-mer is changed. The last group of inequalities is unncessary as we minimize the sum of *y*_*p*_, driving *y*_*p*_ = 0 automatically whenever possible.

#### MinCov

Again, for given *c≤ C*^∗^(*κ, n*), we minimize the number of positions covered by unchanged *k*-mers. A position is covered if it is contained in at least one unchanged *k*-mer. This ILP is an extension of the MinHits formulation, additionally relating the *z*- to the *y*-variables, with a different objective function. We thus need the *x*-, *y*- and *z*-variables.

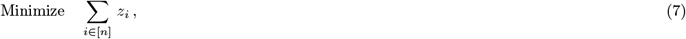

such that (4), (5), (6) hold, and additionally

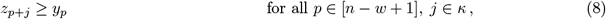

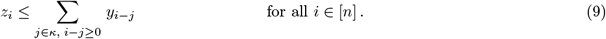

Inequalities (8) force the *z*-variables of all significant positions covered by an unaffected *κ*-mer to 1, whereas (9) forces *z*-variables of positions not covered by any such *κ*-mer to zero. This last group of inequalities is unnecessary because of the direction of the objective function.

#### Solving the ILPs

For modern commercial solvers, the ILPs presented here are of moderate size (see Table 1) and typically solved in a few seconds for a given mask *κ*. The challenge, however, is the large number of (even symmetric) masks of shape (*k, w*), as the ILPs have to be solved for each of the masks. Solution times and optimal masks are given in the Results section.

**Table 1:**
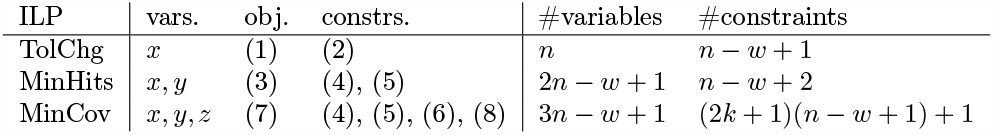
Considered ILPs and their sizes (vars.: used variables, see text; obj.: objective function; constrs.: *necessary* constraints; #: number of).

### 2.4 Strongly Unique, (Weakly) Unique and Non-Unique *κ*-mers

Selecting masks that maximize the MinHits or MinCov criterion gives us error-tolerant primitives in comparison to standard *k*-mers, or minimizers, syncmers, etc. (which are sampled standard *k*-mers).

In most existing sequence comparison tools (read mapping, homology search, etc.), standard or gapped *k*-mers are used as seeds, i.e., short exact matches around which an error-tolerant search or alignment is initiated. With optimized *κ*-mers, we believe that we can propose something more radical: We woud like to showcase the potential of *κ*-mers for one-shot read mapping (without basepair-level alignment).

We classify the *κ*-mers (more precisely, we work with canonical *κ*-mers; hence the restriction to symmetric masks) in a reference genome into the following three types:

**non-unique *κ*-mers** occur at more than one position in the reference,

**weakly unique *κ*-mers** occur at a unique position in the reference, but are closely similar to another *κ*-mer in the reference,

**strongly unique *κ*-mers** occur at a unique position in the reference and have no close similarity to any other *κ*-mer in the reference.

To make the definition concrete and precise, we define “close similarity” as having a Hamming distance of 1.

It is evident that the different types have different implications for mappability: A strongly unique *κ*-mer in a short read can relatively reliably place the read at a unique location in the genome. If several strongly unique *κ*-mers distributed over the read agree on the same genomic location, we can be confident that the read uniquely maps to this location. The same is true, to a lesser degree, for weakly unique *κ*-mers; as they can point to a different location after a single nucleotide change, evidence from several weak *κ*-mers is be required to place the read. Non-unique *κ*-mers can also be helpful (as seeds initiating more detailed search), but cannot be used without further processing. We now face an interesting computational problem.

**Problem 1** *Given a set K of canonical κ-mers, find all pairs* (*x, y*) ∈ *K ×K with x* ≠ *y, such that d*(*x, y*) = 1 *or* 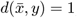, *where d denotes Hamming distance and* 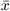 *is the reverse complement of x*.

We previously proposed an implemented a practically efficient solution [21]. Even more efficient approaches may be conceivable, but are not within the scope of this work.

### 2.5 Positional Index of Reference *κ*-mers

Given a reference sequence *s* of length *N* = |*s*| and a mask *κ* of shape (*k, w*), we collect all pairs (*i, x*_*i*_) for *i*∈ [*N* −*w* + 1], where *x*_*i*_ is the canonical *κ*-mer starting at position *i* in *s*. (Invalid *κ*-mers are skipped.) We build a key-value store that maps the (canonical integer representation of the) key *x*_*i*_ to its position *i* if *x*_*i*_ is unique in *s*, and to a special value non-unique if not. Then, the positional information for each *κ*-mer is extended by a strong/weak flag.

To build this positional *κ*-mer index, we have modified our existing gapped *k*-mer counter `hackgap` [22] that uses 3-way bucketed Cuckoo hashing with parallel subtables, to store positional information instead of counts. The hash table can be built from a reference genome in FASTA format.

## 3 Results

We first discuss several interesting masks for typical short reads of length *n* = 100 and their properties. These results are also useful for longer reads, as then they hold for *every* substring of length 100. The appendix contains tables with optimal masks for many shapes (*k, w*) for *n* = 100 and various number of changes *c*∈ { 3, 4, 5, 6, 7} . We then provide information on the solving times of the ILPs from Section 3.2. Section 3.3 contains computational experiments using a challenging set of reads that are on the one hand easily mappable to the t2t reference genome without errors, as the regions they map to are non-repetitive, but on the other hand are hard to map because they contain six substitutions spread over the entire read. Finally, in Section 3.4, we use MiSeq data from the highly variable HLA region (provided to us by the Stefan Morsch Foundation) to evaluate how many reads can be correctly mapped there using different approaches. These results should be understood as a proof-of-concept for the usefulness of optimized masks, and not as claims that we have a better read mapper than existing ones.

### 3.1 Examples of Optimal Masks

For *n* = 100 and (standard) 21-mers, placing 4 changes at positions (20, 41, 62, 83) affects all 21-mers, so the standard *k*-mer mask with *k* = *w* = 21 only tolerates *C*^∗^ = 3 changes. In contrast, the (21, 25)-shaped masks `#####_####_###_####_#####` and `####_#####_###_#####_####` both guarantee at least 8 hits on 100 positions for changes and a minimum of 44 covered positions. Better still, they both tolerate even 5 changes, with at least 3 hits and 33 covered positions. There is no mask with *k* = 21 and 21*≤ w≤* 41 that guarantees more than 8 hits for 4 changes, but the (21, 33)-shaped mask `###_##_# #_###_#_###_# #_##_###` guarantees 55 covered positions or more for 4 changes.

These examples already illustrate the benefit of spaced seeds. Several other interesting masks are collected in Table reftab:masks, which are also used for our computational experiments. Full tables of optimal masks for different shapes (*k, w*) for *c* ∈ *{*3, 4, 5, 6, 7*}* changes on *n* = 100 positions are included in the Appendix.

### 3.2 ILP Solving Times

We used Gurobi 10.0.3 [7] to solve the ILPs from Table 1 on a AMD Ryzen 9 5950X 16-core Processor with hyperthreading (32 logical threads) and 128 GB RAM (which is not needed). Figure 2 shows an overview of solving times for and all 1287 symmetric masks of shape (19, 29) for *c* = 5 changes and sequence length *n* = 100, and all 4368 symmetric masks of shape (25, 35) for *c* = 4 changes (none of theses masks tolerates 5 changes) and *n* = 100. There are a few observations to be made.

**Figure 2:**
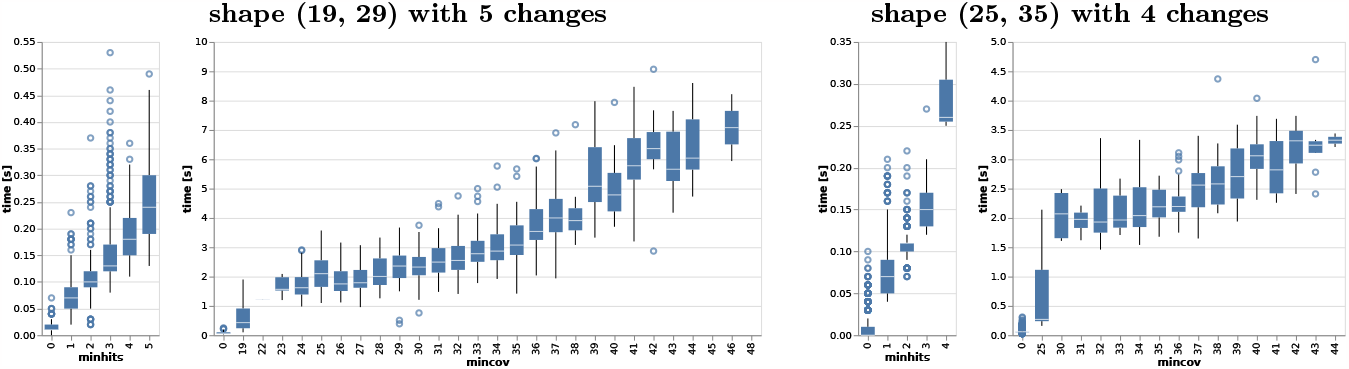
Solving times (y-axis) of the MinCov and MinHits ILP with *n* = 100 for masks of shape (19, 29) with 5 changes (two left plots) and for masks for shape (25, 35) with 4 changes (two right plots). The x-axes show the distinct objective values of the MinHits and MinCov problems. Empty boxplots (e.g. shape (19, 29) with highest MinCov objective value 48) occur if too few masks have this value to produce a meaningful boxplot.

First, solving times are very fast (under half a second) for the MinHits problem and reasonable (under 10 seconds) for the MinCov problem for the chosen parameters. Indeed, the MinHits problem is considerably smaller (Table 1), easier and faster to solve than the MinCov problem.

Second, the solving time depends on the optimal objective value. An objective value of zero (there exists a change distribution for which all *κ*-mers are changed) is found quickly. As a tendency, the better the mask (i.e., the higher the MinHits or MinCov objective value), the harder it is to find the corresponding worst-case change distribution or prove its optimality.

Third, unfortunately, increasing *c* or *n* can have a drastic effect on solving times. This should not be surprising: We are looking for the worst-case placement of *c* changes among *n* positions, for which there are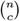 possibilities, where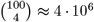 and 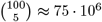. While the time increase between 4 and 5 changes seems moderate in Fig. 2, solving for much larger values of *n* and *c* than provided in the Appendix is currently not feasible in a comprehensive manner. For example, solving the ILPs for the best (19, 29)-mask `###_###_# #_###_# #_###_###` for *n* = 100 and *c* = 5 (guaranteeing 5 hits or 48 covered positions) takes 0.28 seconds for MinHits, and 9.03 seconds for MinCov. Doubling these to *n* = 200 and *c* = 10 increases the time to 41 seconds (*×*146) for MinHits and 5200 (*×*576) seconds for MinCov.

However, we may argue that using *n* = 100 (corresponding to a typical short read length) is sufficient because then the guarantees hold for *every* substring of length 100 of longer reads, and by the pigeon hole principle, if 50 changes are distributed over 1000 positions, there must exist at least one length-100 substring with at most 5 changes. In general, ILP solving times may depend on the decisions taken by the solver and may vary even for the same problem (if randomization is used) and across problems of comparable size and difficulty, and also change from version to version of the solver.

### 3.3 A Challenging Dataset of Reads

We selected 5 million short reads (length *n* = 100) from the autosomes of t2t human reference genome [17]. We introduced *c* = 6 changes into each read at semi-random positions, ensuring that the changes span the whole read. We then used bwa-mem2 v2.2.1 [20], minimap2 v2.26 [11, 12], strobealign v0.11.0 [19] (with default parameters), and the masks A–K from Table 2 to try and map these reads to their position of origin in the t2t genome analysis set (i.e., enriched by an Epstein-Barr virus sequence and a PhiX sequence; see Sec. 3.5). This mapping task is challenging for all of the approaches: Using 6 changes makes it hard for the existing mappers to find a good seed, and none of our masks offers any guarantees for 6 changes either (all MinHits and MinCov objective values are zero). On the other hand, in a sense, the reads were selected to be easy to map because they were taken from repeat-free regions of the genome.

**Table 2:**
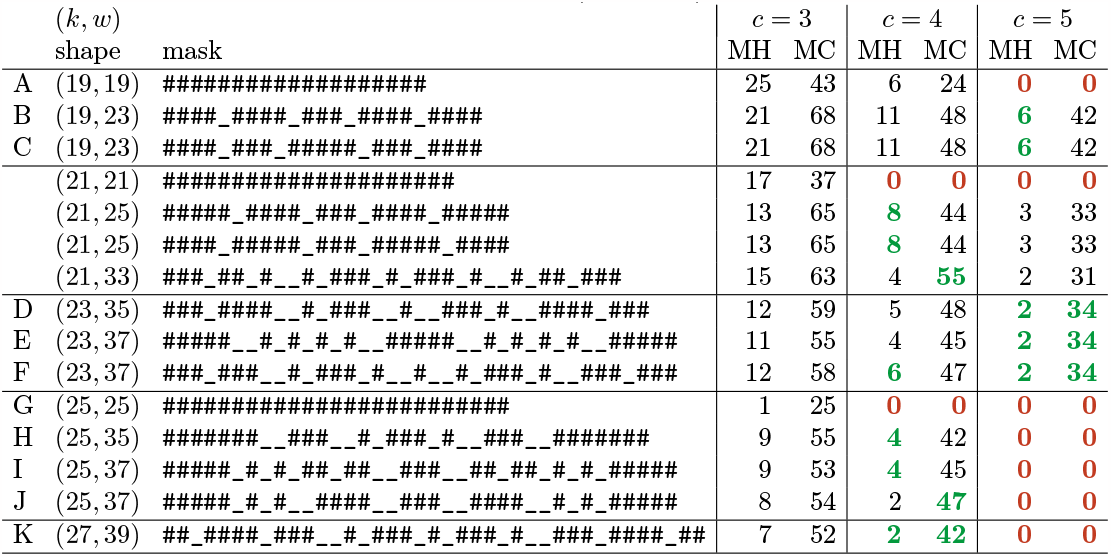
Contiguous *k*-mers vs. well performing masks. Each row shows a mask and its properties for *c*∈ {3, 4, 5} changes in *n* = 100 positions. The label is used to identify the mask in other tables and figures; the masks for *k* = 21 are discussed in Sec. 3.1 and not otherwise used. The shape (*k, w*) is given in addition to the mask for convenience; the remaining columns show MinHits (MH) and MinCov (MC) objective values for different values of *c*. Values of zero are highlighted as **0**; maximal known values for the given *k* (for any *w*) are highlighted in **green**.

We did not create a complete read mapper for gapped *k*-mers, but use a positional index (hash table) as described in Sec. 2.5 for each mask that either stores the unique location (chromosome and position) of a canonical *κ*-mer and whether it is strongly (vs. weakly) unique, or a special value non-unique. Figure 3A contains index statistics and shows at how many positions in the t2t genome we find a strongly unique, weakly unique or non-unique *κ*-mer for the masks A–K from Table 2. Clearly, *k* = 19 (A, B, C) does not yield many strongly unique *κ*-mers, but still an overall high fraction (over 60%) of unique *κ*-mers. We consider it risky to map a read based on a few weak *κ*-mer hits. For *k* = 23 (D, E, F), we already have strongly unique *κ*-mers at over 60% of the genome positions (and a few more positions with weakly unique *κ*-mers). Increasing *k* to 25 (G, H, I, J) or even 27 (K) only slightly increases the fraction of strongly unique *κ*-mers; the fraction of non-unique (“multi”) *κ*-mers stays more or less constant.

**Figure 3:**
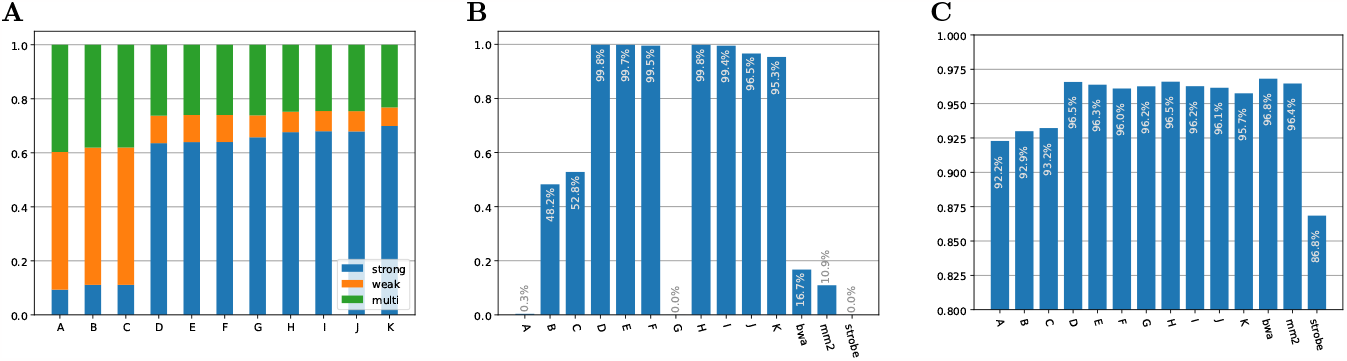
**A:** Fractions of positions in the t2t reference genome where strongly unique (blue), weakly unique (orange) and non-unique (green, “multi”) *κ*-mers start for the masks A–K from Table 2. **B:** Fraction of correctly mapped reads from the challenge dataset (Sec. 3.3). All reads have 6 semi-randomly distributed changes over their 100 positions and are hard to map. Masks A–K refer to Table 2; bwa, mm2 and strobe refer to the established tools bwa2, minimap2 and strobealign. **C:** Fraction of correctly mapped reads from the MiSeq HLA dataset (Sec. 3.4), as in B.

For the alignment tools bwa2, minimap2 and strobealign, a read was considered uniquely mapped if the resulting BAM file contained a unique alignment; it was considered correctly mapped if that unique alignment started at the correct position (with a tolerance of half the read length). For our masks A–K, we queried the genome position of each *κ*-mer in the read and, if unique, subtracted the relative position within the read, to obtain a virtual start position of the read in the genome. We counted (for at most 20 different positions) how many strongly and weakly unique *κ*-mers indicate each position, and aggregate counts to the dominant position (highest count) inside an interval of half the read length. The read is considered uniquely mapped if we find at least two strongly unqiue or four unique *κ*-mers at the dominant position, and that position is the only remaining ones with strong *κ*-mers after aggregation. The read is correctly mapped if the unique position is the correct one in the genome (with a tolerance of half the read length).

Figure 3B shows the fraction of correctly mapped reads for each mask A–K and each tool. The alignment tools struggle with this dataset, because they often do not find a sufficiently long seed (due to the high number of differences) to initiate alignment. In particular, minimap and strobealign suffer because they do not consider all *k*-mers but use a minimizer-based sampling approach. Our contiguous *k*-mer masks (A: 19-mers, G: 25-mers) also perform extremely badly because almost no unchanged 19-mer exists in these reads. In fact, bwa2-mem requires an exact match with a minimum length of 19 to initiate further alignment. As it also uses non-unique 19-mers (and investigates up to 1000 positions), it is more successful that our approach that only uses (ideally strongly) unique 19-mers. The change-tolerant gapped 19-mers (B, C) perform much better than the contiguous 19-mer (A), but still not very well. The limiting factor here is the unavailability of sufficiently many strongly unique gapped 19-mers (Fig. 3A). Our masks with *k* = 23 perform well (D, E, F), even though they guarantee only 2 hits for 5 changes, and no hits for 6 changes. Because the change distribution is semi-random, we typically do not see the masks’ worst-case distributions and therefore usually get the required two strongly unique *k*-mer hits even for 6 changes. The same is true for the gapped 25-mer masks that guarantee 4 hits for 4 changes (H, I), but the remaining masks (J, K) that guarantee only 2 hits for 4 changes map slightly fewer reads.

### 3.4 Mappability of MiSeq Data to the Human HLA Region

The Stefan Morsch Foundation (Birkenfeld, Germany) offers free HLA typing (and sequencing) for potential stem cell donors. The data is not available for general public use. We use collected reads from targeted HLA sequencing (MiSeq paired-end, but reads were used as single-end) from 79 samples, with an average of 125 000 read pairs (250 000 reads) per sample. In principle, all reads should map to the 11 Mbp HLA region on human chromosome 6. Similar to the challenge data, we consider a read correctly mapped if it is mapped to a unique location and that location is within or close to the HLA region (1 Mbp tolerance).

Figure 3C shows that most approaches do this successfully, mapping over 96% of the reads to the correct region. Both bwa2-mem and minimap2 find enough exact contiguous seeds, but strobealign is less successful (note that the scale on the y-axis starts at 80% to highlight the small differences). As before, our 19-mer masks suffer from the scarcity of (strongly) unique 19-mers, but the 23-mer and 25-mer masks perform well. Two masks (D, H; both 96.5%) are competitive with bwa2-mem (96.6%), although we never perform alignment and solely rely on *κ*-mer hits.

### 3.5 Software and Data

With this article, we provide the following tools^1^, further explained in the accompanying README file: (1.) a Python program (that calls the Gurobi optimizer [7], for which a (free academic) license is required) to compute the worst-case change distributions of a mask or a set of masks of the same shape, (2.) a just-in-time compiled Python program to index the *κ*-mers of a genome in FASTA format (`gapmap index`), which is a modification of our gapped *k*-mer counter hackgap [22], (3.) a just-in-time compiled Python program to map a FASTQ file of reads using a pre-computed *κ*-mer index (gapmap map) We furthermore provide the following data: (4.) a list of worst-case optimal masks for different shapes (*k, w*) and number of changes *c* for *n* = 100 in Appendix A, (5.) our modified t2t reference genome (“analysis set”) consisting of the t2t genome [17], an Epstein-Barr virus (EBV) sequence and a PhiX sequence,^2^, (6.) the challenge dataset of Sec. 3.3 of 100-bp reads with 6 changes.^3^ The FASTQ headers contain the true origins and the positions of the introduced changes.

## 4 Discussion and Conclusion

We have opened a new chapter in spaced seed design. Instead of considering the probability or guarantee of at least one hit, we use a combinatorial approach and explicitly compute the worst-case distribution of *c* changes (substitutions) among *n* positions to minimize the number of hits or the number of positions covered by hits. The integer linear program formulations we use for this task are reasonably efficient for *n* = 100 and allow exhaustive enumeration of all symmetric masks for a wide range of shapes (*k, w*); but they show their limits for longer sequences: Already for *n* = 200, solving times for a single mask prohibit evaluating most shapes of interest. Further engineering, such as the inclusion of starting and improvement heuristics, and exploitation of the regular constraint structure, may lead to substantial performance improvements.

We focus on shapes (*k, w*) for which most *κ*-mers in a mammalian genome can be expected to be unique or even strongly unique with the idea that a few strongly unique *κ*-mer matches suffice to map a read to a specific position of the reference genome. As demonstrated, this works in the human genome for *k ≥* 23, for which many strongly unique *k*-mers exist. This is different from previous work that focused on small *k*, and used matching *κ*-mers as seeds to initiate alignments at potentially many different positions where the *κ*-mer occurs. The approach based on strongly unique *κ*-mers can be orders of magnitude faster and at the same time highly reliable, as shown by our challenge dataset and the HLA dataset. Thus, strongly unique worst-case-optimal spaced seeds provide an orthogonal method against reference bias besides graph genomes.

We found that good worst-case masks are rare, so we cannot expect to find a good mask randomly. Also, which mask is optimal may differ for different values of *c* (for the same *k, w* and *n*). This suggests using a combination of several masks, which would also increase the chance of seeing more strongly unique *κ*-mers (across all used masks). The problem then becomes finding an optimal combination of masks (perhaps even of different shapes). This has already been considered within the probabilistic approach, but an exhaustive enumeration of all combinations is not feasible, so heuristics have been proposed, which may be adapted to worst-case mask design in the future.

A drawback of spaced seeds is their rigidness over their width *w*, i.e., they tolerate substitutions, but not insertion- or deletion-type differences. There is a trade-off: A longer width may lead to more hits or higher coverage, but also offers less tolerance against insertions or deletions. While strobemers were developed with indel tolerance in mind, they did not perform well in the adversarial settings considered here. From our point of view, it remains a challenging open problem to alternative general error-tolerant types of seeds.

There are other open questions as well: We are currently unable to predict the performance of a mask (e.g., from substrings, motifs, etc.). It would be interesting to use machine learning approaches to predict which masks have the potential to yield high MinHits or MinCov values to avoid running all ILPs. Efficiently finding all pairs of *κ*-mers with Hamming distance 1 in a large set *X* of *κ*-mers is an interesting algorithmic problem which so far has received little attention, and while we have a reasonably efficient working solution [21], better approaches may exist.

## A Appendix: Tables of Optimal Masks for 100 Positions

The following tables show optimal masks for *n* = 100 positions and different numbers of changes *c* ∈ { 3, 4, 5, 6, 7} . There is one long table for every value of *c*. Each table is sorted by shape (*k, w*) for reasonable choices of shapes.

A mask is included in the table if it is optimal for its shape, i.e., either it has maximal MinHits value or maximal MinCov value among all masks of the same shape. The corresponding values are given in columns **hits** and **covpos**, respectively. A value is highlighted in **green** if the value is maximal among all evaluated masks with the same *k*, irrespective of *w*. It is possible that even better masks exist for larger *w* than evaluated so far. An additional column **tol+** indicates whether the mask tolerates more changes (and how many) than used for the current table.

Values in parantheses, if present, indicate, in the **hits** column, the guaranteed number of covered positions if the change distribution achieves the worst-case number of hits; and in the **covpos** column, the guaranteed number of hits if the change distribution achieves the worst-case number of covered positions.

**Example**. Consider the following two rows from the table for *c* = 3 changes and shape (*k, w*) = (21, 27).

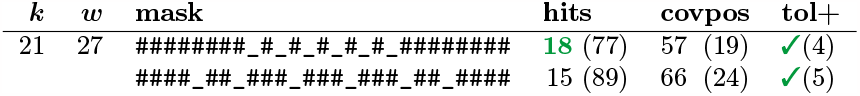

The first (upper) mask guarantees 18 hits (unchanged *κ*-mers) for all possible distributions of 3 changes over 100 positions. This is maximal for this shape, and furthermore, because **18** is highlighted, it is maximal for *k* = 21 over all choices of *w ≥ k*. It also guarantees to cover 57 of the 100 positions with unchanged *κ*-mers, but this is not maximal for this shape. Indeed, the second mask guarantees 66 covered positions, which is maximal for shape (21, 27), but not maximal for *k* = 21 overall, because there is a mask of shape (21, 41) that guarantees even 78 covered positions. The second mask also guarantees 15 hits, which is not maximal.

Interestingly, the first mask guarantees more hits for 3 changes and tolerates 4 changes, but not 5 changes (i.e. there is a distribution of 5 changes over 100 positions that changes all *κ*-mers), while the second mask guanrantees fewer hits for 3 changes but tolerates up to 5 changes.

Let us now focus on the values in parentheses for the second mask, 15 (89) and 66 (24). They show that the worst-case change distributions to achieve MinHits or MinCov are different. More precisely, any change distribution that yields only 15 hits guarantees at least 89 covered positions, while any change distribution that yields only 66 covered positions guarantees at least 24 hits.

These values have been computed by a modified ILP that minimizes the covered positions (resp. hits) while constraining the hits (resp. covered positions) to be equal to their worst-case (minimal) value obtained from the original ILP.

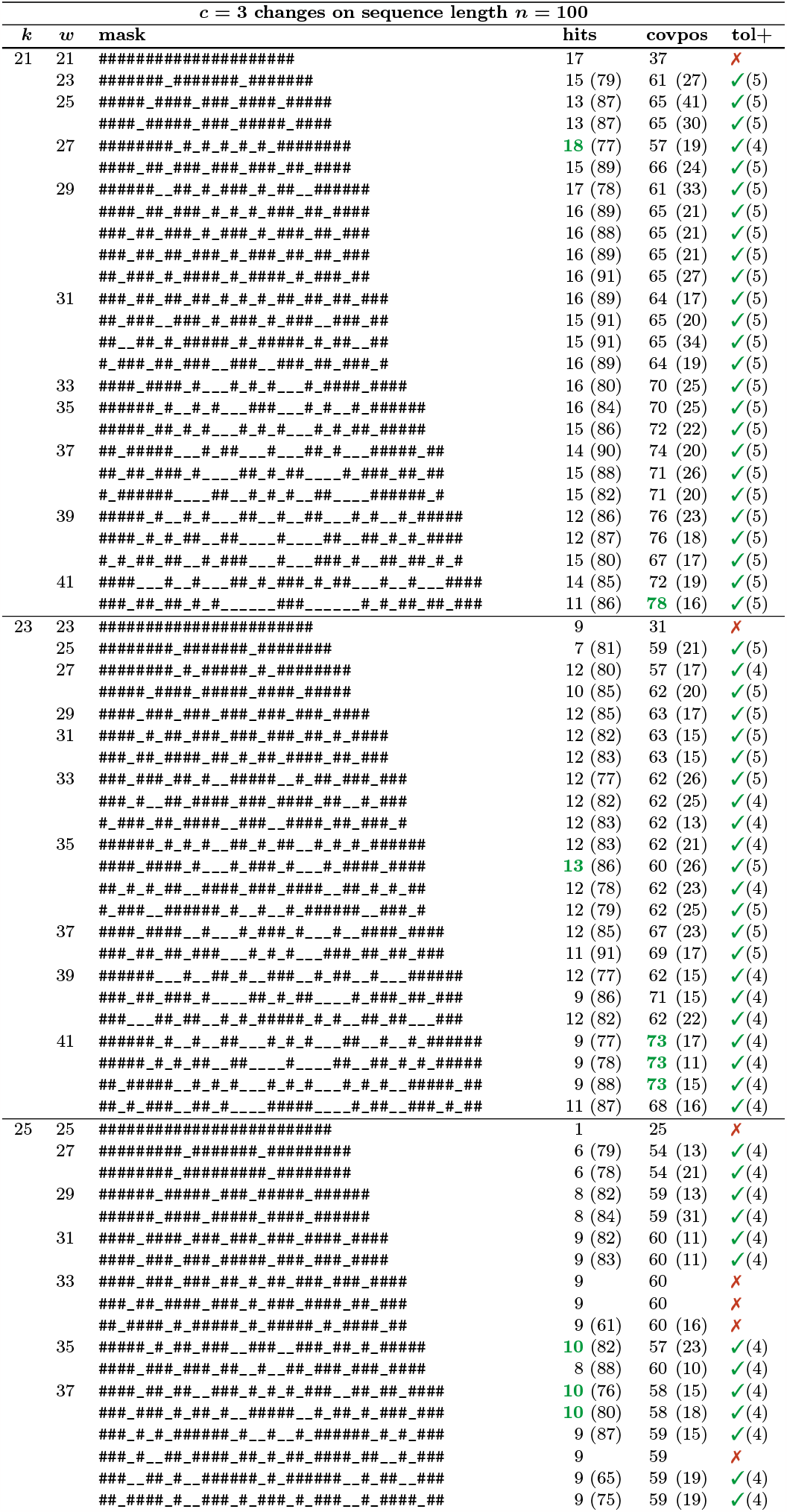
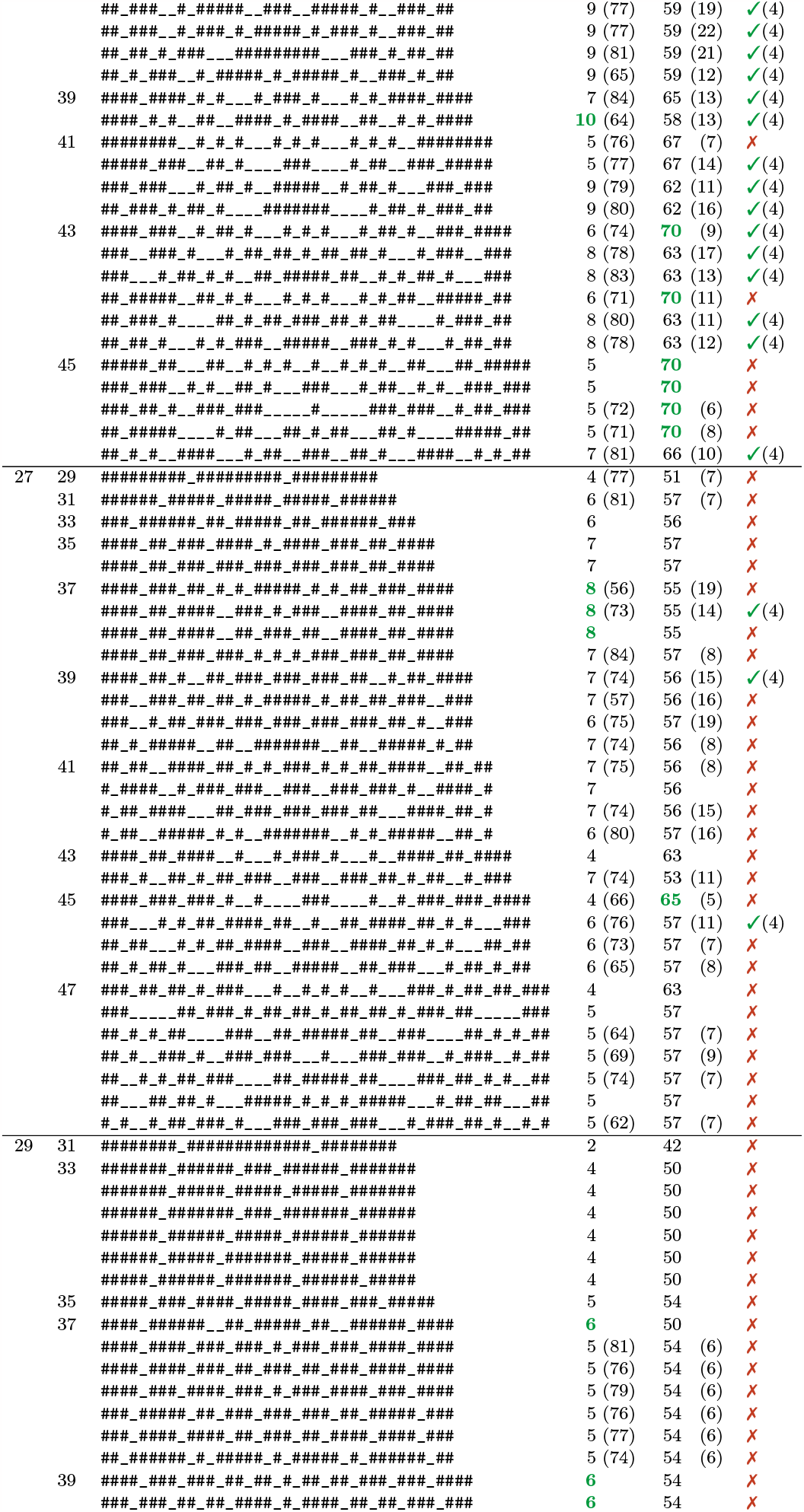
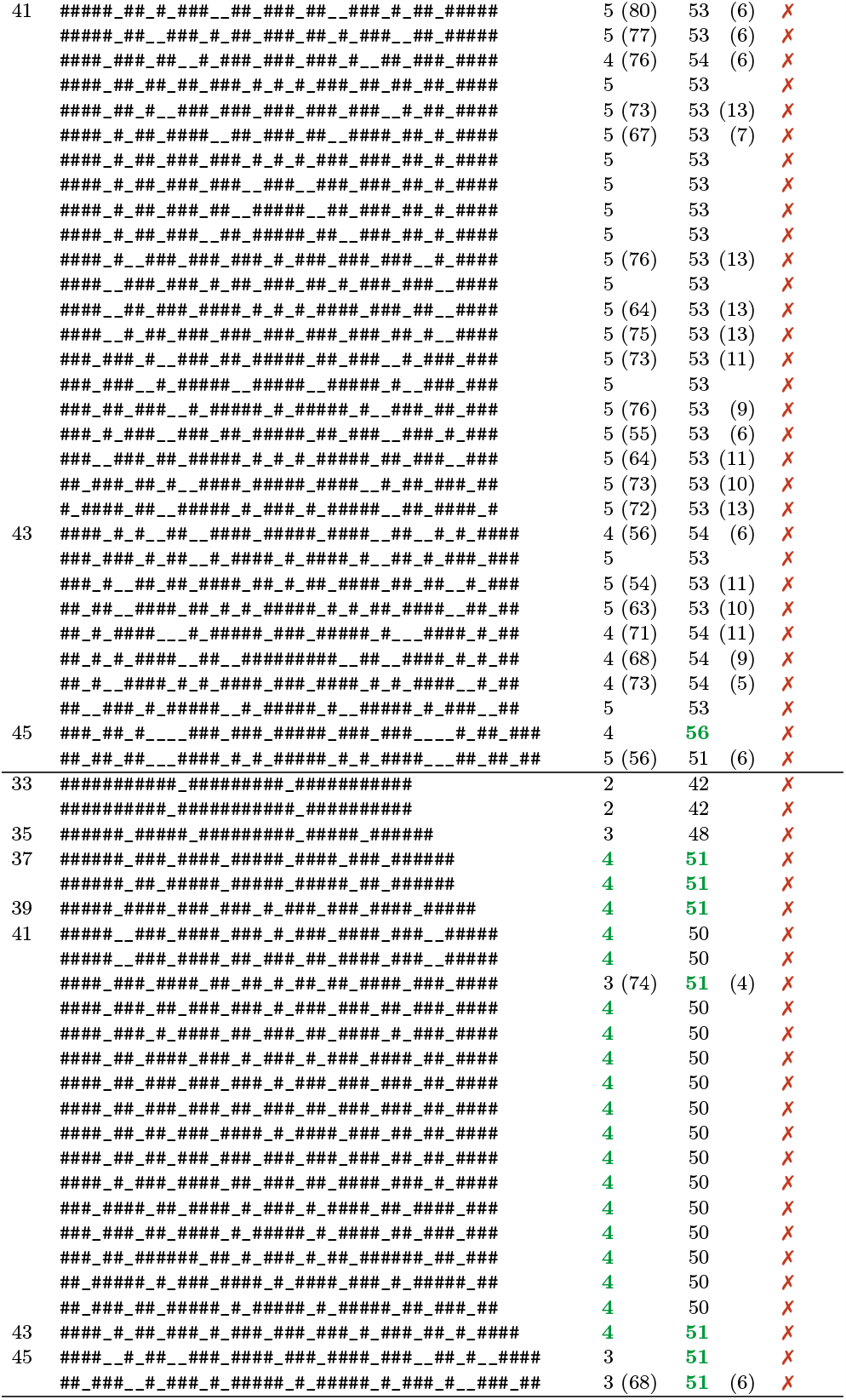

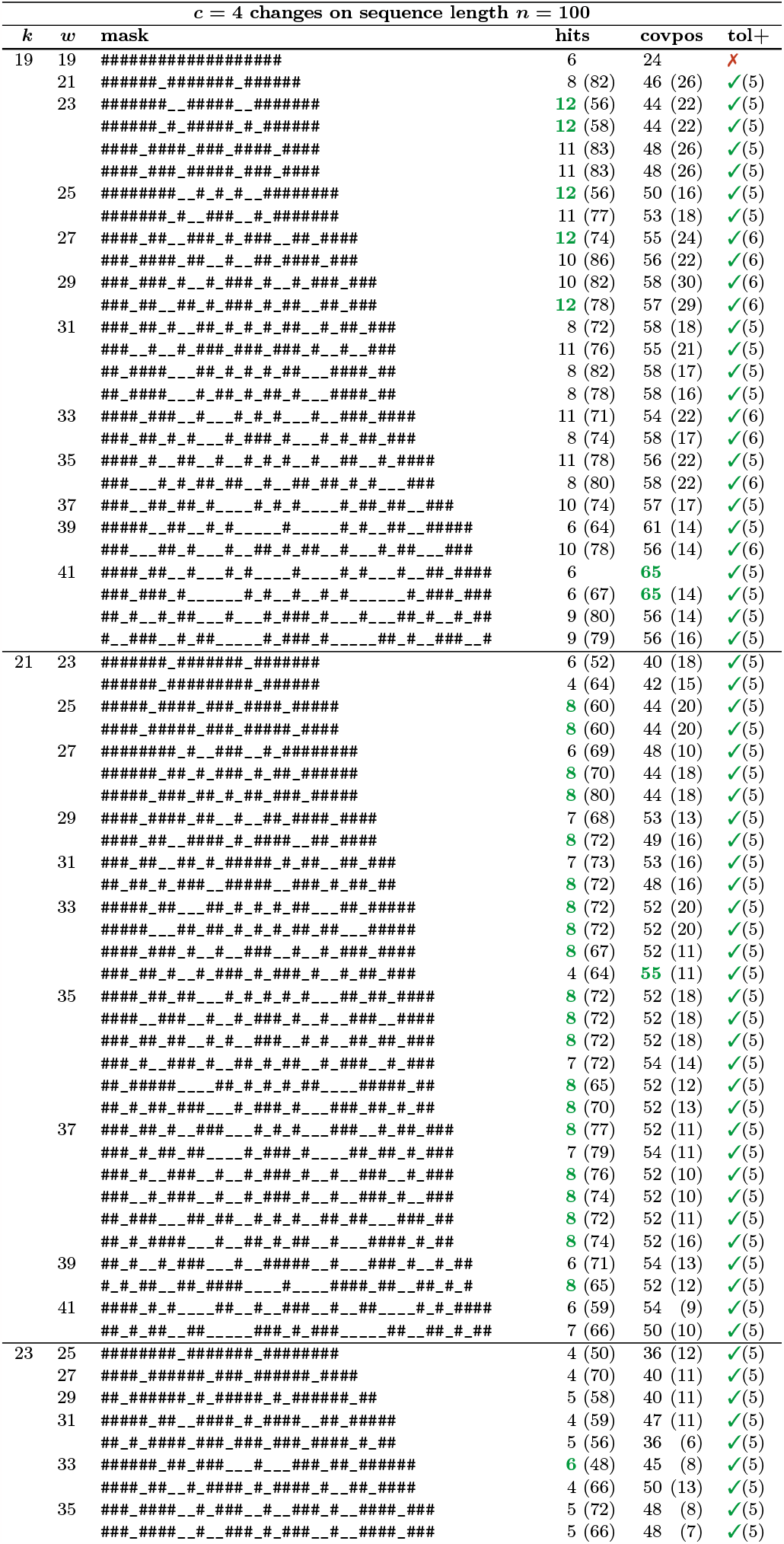
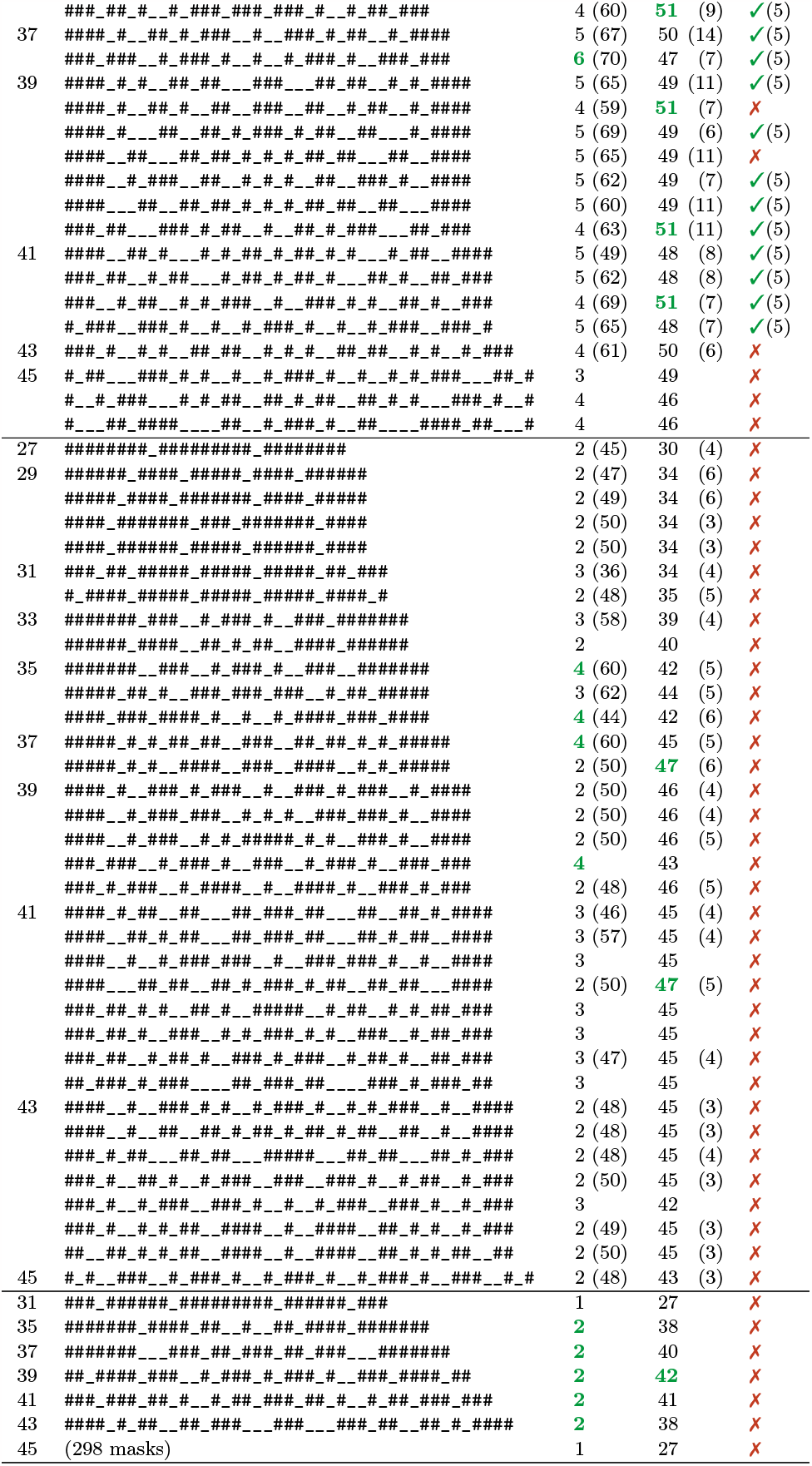

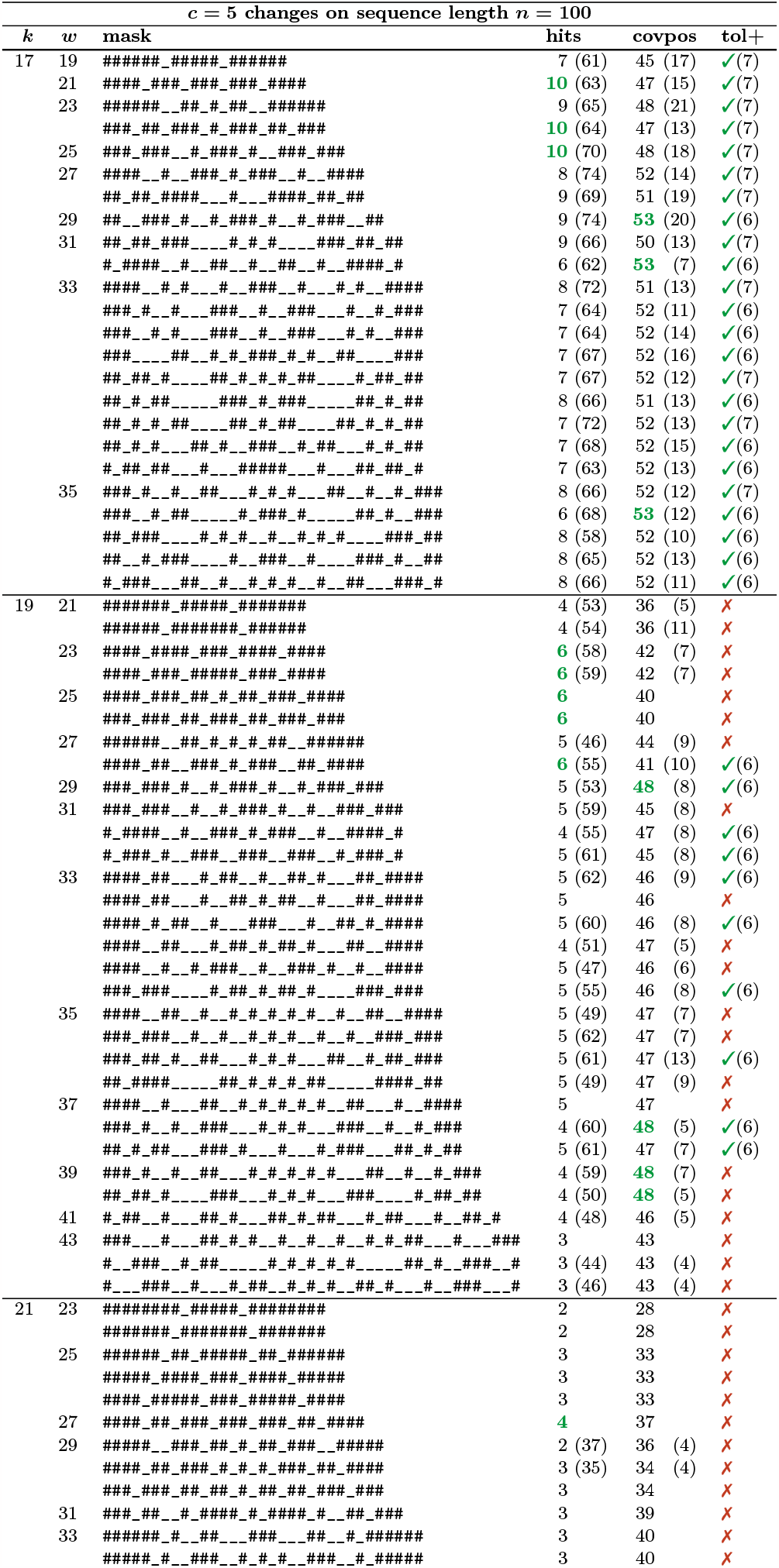
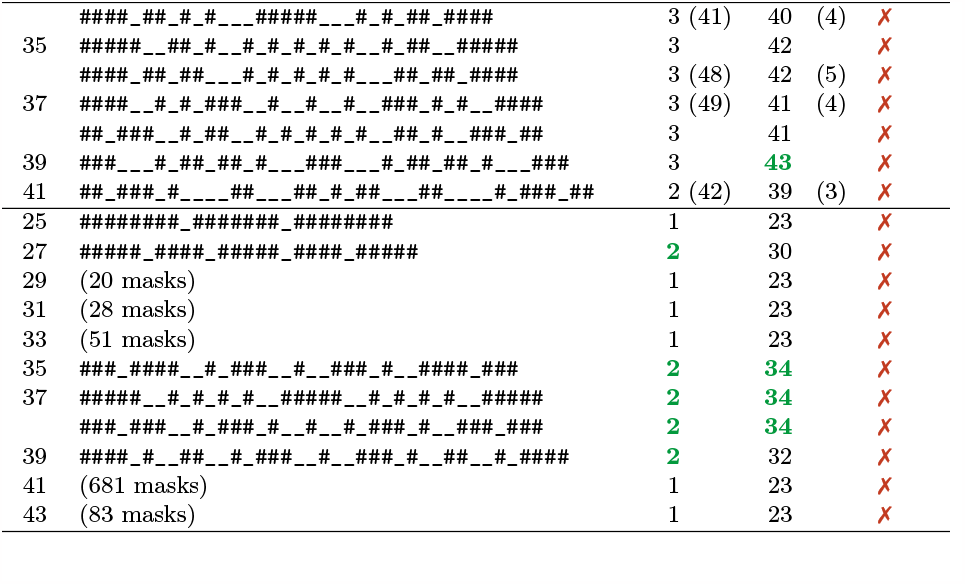

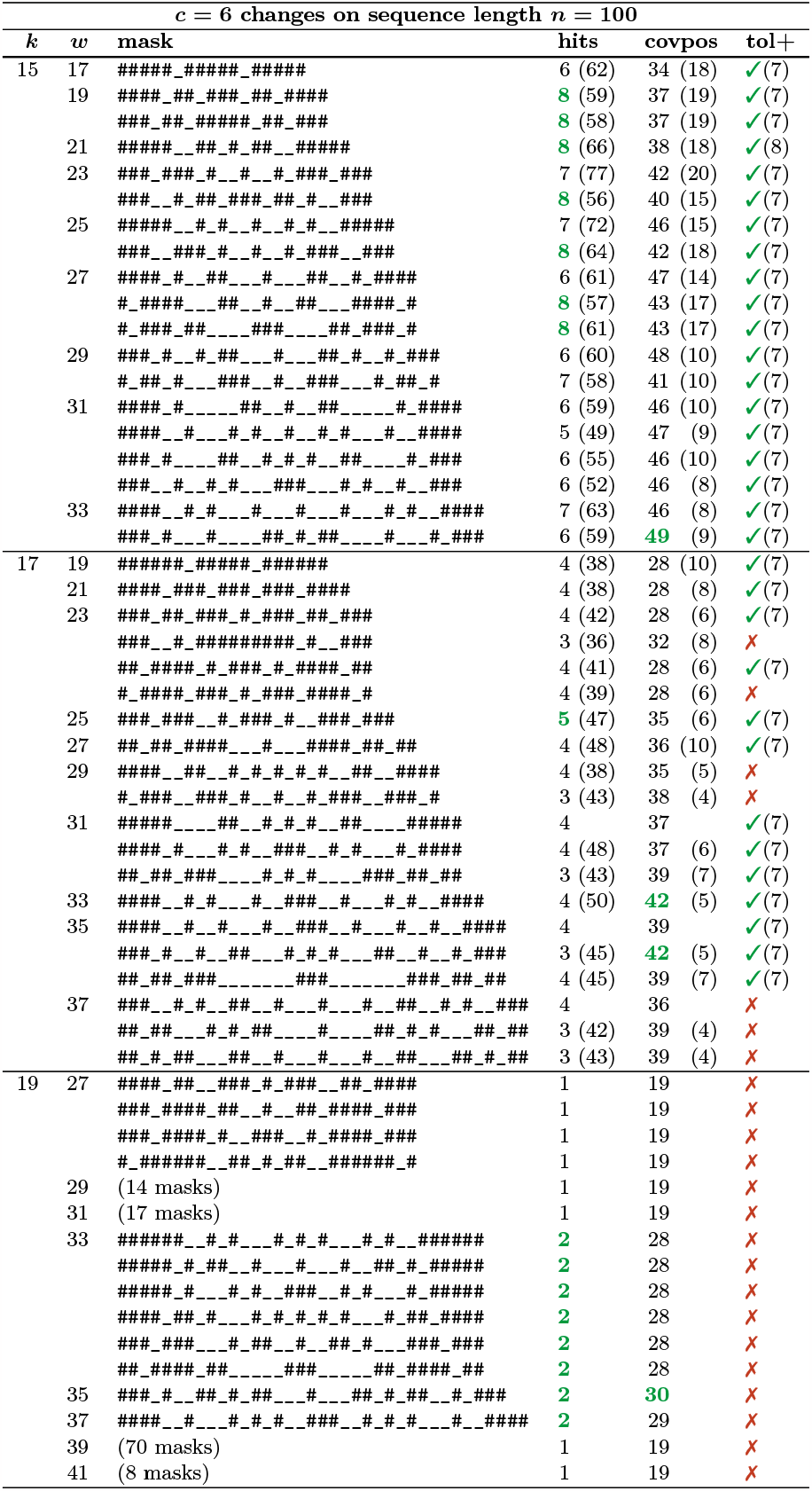
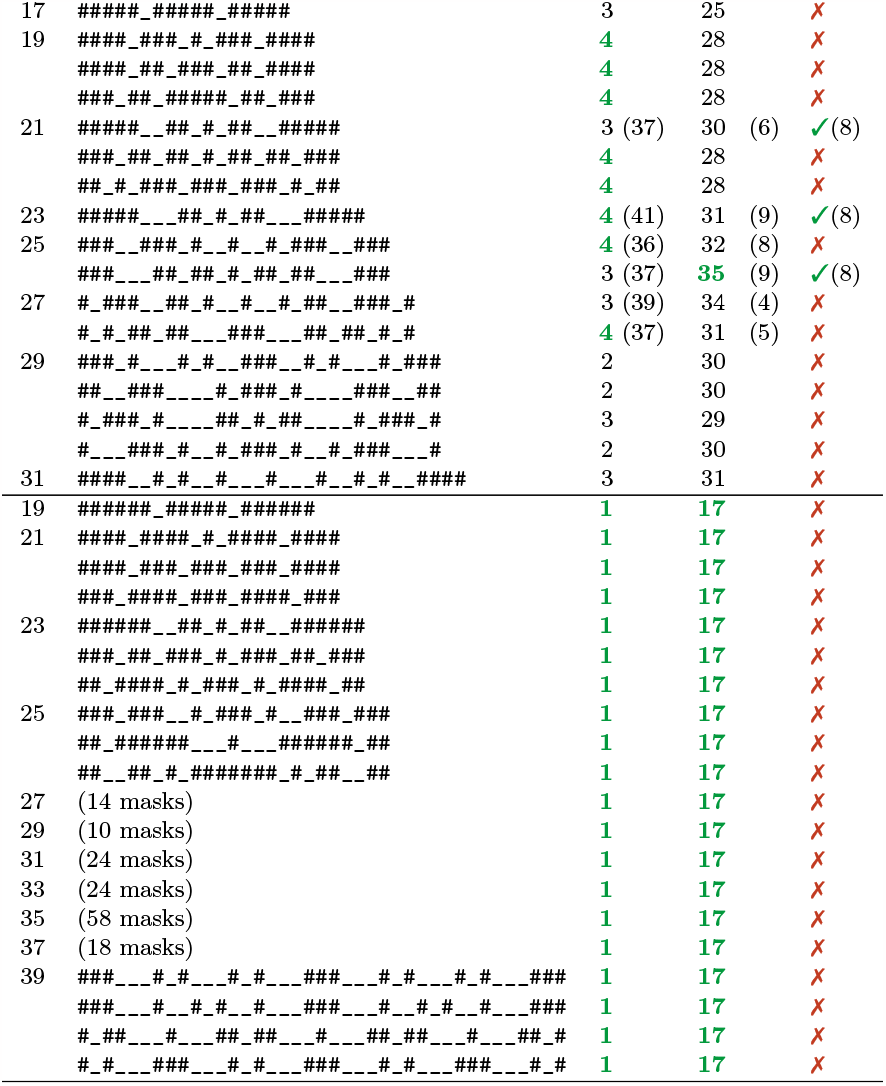

## Footnotes

1 https://gitlab.com/rahmannlab/seed-optimization

2 https://kingsx.cs.uni-saarland.de/index.php/s/CcggyrNZWpFoZWj

3 https://kingsx.cs.uni-saarland.de/index.php/s/mA9cXMYcgHsZXFb

